# *Daphnia magna* egg piRNA cluster expression profiles change as mothers age

**DOI:** 10.1101/2021.11.05.467411

**Authors:** Jack Hearn, Tom J. Little

## Abstract

PiRNAs prevent transposable elements wreaking havoc on the germline genome. Changes in piRNA expression over the lifetime of an individual may impact on ageing through continued suppression, or release, of transposable element expression. We identified piRNA producing clusters in the genome of *Daphnia magna* by a combination of bioinformatic methods, and then contrasted their expression between parthenogenetically produced eggs representing maternally-deposited germline piRNAs of young (having their 1^st^ clutch) and old (having their 5^th^ clutch) mothers. Results from eggs were compared to cluster expression in three generations of adults. As for other arthropods, *D. magna* encodes long uni-directionally transcribed non-coding RNAs which consist of transposable element fragments which account for most piRNAs expressed. Egg tissues showed extensive differences between clutches from young mothers and those from old mothers, with 578 and 686 piRNA clusters upregulated, respectively, although most log fold-change differences for significant clusters were modest. When considering only highly expressed clusters, there was a bias towards 1^st^ clutch eggs at 41 upregulated versus eight clusters in the eggs from older mothers. F_0_ generation differences between young and old mothers were fewer than eggs, as 179 clusters were up-regulated in young versus 170 old mothers. This dropped to 31 versus 22 piRNA clusters when comparing adults in the F_1_ generation, and no differences were detected in the F_3_ generation. These patterns were similar to that observed for *D. magna* micro-RNA expression. Little overlap in differentially expressed clusters was found between adults containing mixed somatic and germline (ovary) tissues and germ-line representing eggs. A cluster encompassing a Tudor domain containing gene important in the piRNA pathway was upregulated in the eggs from old mothers. We hypothesise that regulation of this gene this could form part of a feedback loop that reduces piRNA pathway activity explaining the reduced number of highly-expressed clusters in eggs from old mothers.

**Author Summary:** Small RNAs shorter than 200 nucleotides often function by targeting RNAs with complementary nucleotide sequences for destruction. A subset of small RNAs, the Piwi-interacting RNAs or piRNAs are best known for silencing sequences of DNA that can jump between locations in the genome which can compromise the integrity of genomes. This protects offspring from sterility and other undesirable effects, hence piRNAs have been termed ‘guardians of the genome’. PiRNAs have several characteristics, such as a characteristic length and origin in genomic graveyards of junk DNA, that can be used to distinguish them from other small RNAs. Here, we used a combination of computational approaches to identify piRNA-producing clusters in the genome of the water flea *Daphnia magna*. We then contrasted expression of these clusters between genetically identical adults of different ages, their eggs, adult offspring and great-granddaughters. Adults and their eggs were markedly different in cluster expression by age, which was mostly lost by adulthood in offspring and not seen at all in great-granddaughters. By taking an innovative approach that can be applied to similar datasets of diverse organisms we have shown that piRNA expression, and therefore, stability of the genome can change with age.

## Introduction

### Roles of piRNAs and their production

Piwi-interacting RNAs (piRNAs) are 21-35 nucleotide long small RNAs that are maternally deposited into oocytes to provide immunity to complementary transposable elements (TEs) by suppressing new insertions into the germline. This protects the developing embryo from transposon-mediated illegitimate recombination, double-stranded breaks, and disruptive insertions into coding sequences promoters which can cause aberrant gene expression [1,2]. Although efficient against recognised TEs of maternal origin, piRNAs are less effective against TEs of paternal origin [2]. In arthropods, although research has focussed on germline piRNAs, somatically expressed piRNAs are an ancestral mechanism of protection against TEs [3]. This transposon suppression role may even represent a deeper ancestral trait of bilaterians [4,5]. Additional functions of piRNAs include regulating gene expression, protecting against viruses and telomere maintenance [2,6], and, exceptionally, piRNAs act as master-regulators of sex-determination [7] in silkworms.

PiRNAs originate from long precursors RNAs in arthropods and mammals [6], and are first transcribed and then further processed in the cytoplasm into mature piRNAs [6]. These piRNA-producing clusters contain contiguous remnants and nested fragments of transposons consigned to genomic ‘graveyards’ [2,6]. They are transcribed in a single direction in most arthropods surveyed to date, with *Drosophila* species also having evolved a unique dual-strand stranded system expressed by germline cells [8,9]. After transcription, piRNA cluster transcripts enter either the ping-pong cycle or phased piRNA pathways which both occur in the cytoplasm [2,6]. Phased piRNA production can occur in germline and somatic cells and occurs when Piwi (P-element induced wimpy testis) is loaded at the 5’ end of piRNA precursors and cleaved by the Zucchini protein [10]. The process is then repeated in a step-by-step process through Piwi loading of newly created 5’ ends to produce distinct piRNAs along the precursor RNA [11]. The ping-pong cycle occurs in germline cells when a sense orientated piRNA guides the Argonaute 3 (Ago3) protein to a complementary cluster RNA and cleaves it [11]. Once Ago3 and a piRNA are bound, the Aubergine (Aub) protein binds to the 5’ end of the Ago3-piRNA targeted mRNA cleavage products and slices it into a mature anti-sense piRNA which recognises and cuts complementary RNAs such as TE mRNA. Ago3 recognises the 5’ ends of these cleaved RNAs thus propagating the ping-pong loop which ultimately results in the post-transcriptional silencing of transposons [11]. Maternal deposition of the Aub protein and associated piRNAs into oocytes initiates the ping-pong cycle intergenerationally in *Drosophila* [1,12].

Together, the ping-pong loop and phased piRNA pathways create a diverse population of piRNAs optimised for their RNA silencing role [2]. In addition to post-transcriptional suppression of targets, both pathways also can direct transcriptional silencing in the nucleus [2]. PiRNAs guide Ago3 to complementary transposable element insertion sites in the nucleus and promote H3K9me3 modification of histone tails which leads to heterochromatin formation and ultimately silencing of the locus [2,6,11]. The characteristic size of piRNAs is determined by loading of intermediate piRNAs into Piwi and Aub proteins. Due to intrinsic preferences of Piwi proteins in each pathway, phased pathway piRNAs have a bias for Uridine at the 1^st^ base (1U) and ping-pong cycle piRNAs for Adenine at the 10^th^ position (10A) with no associated 1^st^ base bias [6]. Together, piRNA length distributions and base position biases are useful characteristics for classifying piRNAs versus other sRNA species.

The association between Piwi and piRNAs is weaker than that for miRNAs with Ago1 [13]. PiRNAs therefore require a longer region of matching with their target RNA than the seven base pairs in miRNAs to form a stable association. As a result, piRNA target recognition is more selective than for miRNAs, but beyond an essential seed-matching region, mismatches between piRNA and their targets is tolerated. This can lead to evolutionarily conserved piRNA-target interactions despite the accumulation of mutations in transposable element sequences over time [13].

### Ageing and the piRNA pathway in arthropods

Ageing is associated with increased expression of transposable elements in a variety of animals due to progressive genomic dysregulation [14,15], with age-related phenotypes resulting from negative effects of transposition on cell and genome integrity [14]. Among arthropods, lifespan has been shown to increase in *D. melanogaster* through the application of reverse transcriptase inhibitors which reduced TE activity [14,16]. The piRNA pathway was observed to differ intergenerationally with age in *D. melanogaster*, as egg chambers of older mothers were upregulated for piRNA pathway genes (27 of 31 tested) [17]. PiRNA pathway genes have also been implicated in somatic ageing of the termite *Macrotermes bellicosus* [18]. The heads of older, worker caste termites had lower expression of four such genes and higher expression of several hundred TEs versus younger workers suggestive of reduced suppressive activity [14,18]. By contrast much longer-lived termite kings and queens maintain stable TE and gene expression levels throughout lifetimes [18]. In line with termite workers, increased TE activity with age occurs in *Drosophila* somatic tissues [19–21], albeit with the caveat that TE insertion rates are prone to overestimation in sequencing data [22]. It has been hypothesised that that age-related misexpression of TEs is restricted to somatic tissues due to continued efficient policing of the germline by the piRNA response with age [17,18,23].

### The piRNA pathway in crustaceans

The piRNA pathway is likely to underly the evolution of obligately parthenogenetic reproduction from cyclical parthenogenesis in the cladoceran crustacean *Daphnia pulex* [24]. A transposon insertion upstream of the meiotic cohesion factor *Rec8* in *D. pulex* was found to correlate perfectly with obligate parthenogenesis in effected strains [24]. The piRNA pathway is hypothesised to silence the TE-inserted copy of *Rec8* and several other wildtype paralogs of *Rec8* through sequence homology to the generated piRNAs [24]. Little further is known of the presence and action of piRNAs in *Daphnia* or generally across the crustacea, nor what impact a cyclically-parthenogenetic lifecycle has on piRNA dynamics. *Daphnia magna* and *Daphnia pulex* each encode seven essential piRNA pathway Piwi-like genes in their genomes [25,26], alongside three Argonaute genes in *D. magna* to two in *D. pulex* [25]. Other crustaceans are likely to encode piRNA pathway genes, with evidence for piRNA expression and piRNA pathway genes in *Triops cancriformis* (tadpole shrimps, class Branchiopoda, order Notostraca) [27]. Seven copies of Piwi in *Daphnia* species represents a gene expansion versus other arthropods, which is perhaps associated with sub-functionalisation across Piwi copies to the soma and splitting of germline roles between meiosis and parthenogenesis of cyclically parthenogenetic species [28]. A similar association between expanded Piwi-like genes and reproductive plasticity was observed in the genome of the pea aphid *Acyrthosiphon pisum* [29]. In addition to gene expansion, genes of the piRNA pathway exhibit strong signals of positive selection and expansion in *Drosophila* species, probably in response to repeated transposable element invasions and in antiviral defence [30,31].

### *Daphnia magna* piRNA responses to ageing

Prior research has established that micro-RNA (miRNA) expression and DNA methylation status respond to ageing and caloric restriction in *D. magna* [32–34], with caloric restriction resulting in pervasive effects on gene expression [35]. The miRNA and DNA methylation profile of mothers changed with age in parthenogenetically reproducing individuals [32,34]. The eggs of old mothers also showed a different miRNA profile from the eggs of young mothers, but this difference was greatly reduced once eggs hatched and grew to adulthood, and was not evident in great granddaughters [32]. Here, we take advantage of that pre-existing resource to rigorously identify piRNAs in eggs and adult tissues of *D. magna* for the first time, which we used to predict putatively piRNA producing loci along the *D. magna* genome. By quantifying clusters in eggs from primiparous and multiparous (specifically, having their 5^th^ clutch) mothers, we tested for age-related changes in piRNA expression in eggs and their maternal and descendent generation adult *Daphnia*. There were extensive differences between eggs from young and old mothers, with little overlap in differentially expressed clusters in their mothers, akin to the pattern observed in miRNAs. PiRNA expression is therefore regulated to some extent by ageing-related processes in *D. magna*.

## Results

### piRNA cluster identification

Quality and length filtering of reads had the largest effect on egg small RNA libraries with 38-72% of data remaining after this step, removal of miRNA, other non-coding RNAs and piRNA classification had a lesser effect (Table S1). After all filtering steps 35-64% of original reads were retained, and absolute numbers of reads per egg library ranged from 5.2 to 10.5 million. Results for adult piRNA libraries were similar with one outlier (replicate Y1A_F0, Table S1). There were 15,719 ShortStack clusters after aligning egg and adult libraries (Table 1), of these 4,747 had a significant (adjusted p-value < 0.05) PingPongPro predicted ping-pong cycle signature. ProTRAC predicted 19 piRNA producing loci each for egg and adult libraries analysed separately (File S1). On combining egg and adult proTRAC results 22 unique piRNA producing loci remained, all of which were transcribed uni-directionally. After merging ShortStack and proTRAC predicted loci, removal of clusters with an average read length of 24 or less, and clusters overlapping other species of RNA in the genome annotation 4,606 putative piRNA loci remained for expression-based analyses, including 20 proTRAC clusters. These 4,606 clusters had an average aligned read length of 26.27 bp (standard deviation, ±0.53). The proTRAC predicted clusters spanned 2.1% of all filtered putatively piRNA producing loci (466,865/22,296,837 bp), whilst accounting for on an average of 47% and 56% total expression (as proportion of normalised counts) in eggs and adults respectively (Table S2). Cluster lengths varied from 31 to 91,046 bp and were present on all 10 linkage groups of the *D. magna* genome.

**Table 1.** Numbers of piRNA clusters predicted and after filtering for input to differential expression analysis. The final row gives the number of clusters input to differential expression analysis.

| Predictions | Number |
| --- | --- |
| ShortStack predicted clusters. | 15,719 |
| ShortStack clusters containing PingPongPro predicted transposons. | 4,747 |
| Adult proTRAC cluster predictions. | 19 |
| Egg proTRAC cluster predictions. | 19 |
| Merged proTRAC cluster predictions. | 23 |
| PingPongPro and proTRAC merged predictions. | 4,732 |
| Average read length filtered clusters. | 4,653 |
| Final count after clusters overlapping other RNA species were removed. | 4,607 |

### Egg versus adult piRNA cluster expression

Many clusters had large expression differences between egg replicates combined and adult replicates combined (F_0_ + F_1_ + F_3_ generations). For egg transcripts per million (TPM) per cluster averaged across replicates, 33 had a TPM of 1000 or more greater than adult averages. Conversely, 37 clusters had a TPM greater than 1000 or more in the average of adult replicates relative to eggs (Table S3). Three of the clusters upregulated in eggs had much greater differences in TPM (13,354-20,060 TPM) versus the remainder (1,059-6,130 TPM). Two of these most upregulated clusters were proTRAC predicted piRNA producing loci occurring on the negative strand. NC_046175.1:10253003-10266628 at 13.6 kb long encompasses a long non-coding RNA (lncRNA) which act as precursors to piRNAs flanked by two protein coding genes. Most piRNAs aligned in the region within the lncRNA and were biased toward primary piRNAs, with 96% of 1^st^ read positions being uracil and 27% adenine at the 10^th^ position. The second cluster had a similarly biased 1U to 10A ratio (88% to 22%). For this cluster, no overlapping lncRNA has been annotated and most reads align across and downstream of an uncharacterised gene ‘LOC116926960’. The third cluster not within a proTRAC prediction was very short at 35 bp long and did not overlap a feature, the closest gene-body was ∼3 kb downstream of the cluster and encodes a lncRNA locus. Seven adult clusters had a TPM difference greater than 10,000 in favour of adults versus eggs. Of these, six are proTRAC clusters and the remaining locus was annotated with retro-element domains by eggNOG. Further to these high-expression clusters, two consecutive clusters separated by only 2.8 kb had very low average TPMs (< 10) in eggs versus adults (> 1000). Both clusters overlap a single gene (LOC116928002) which encodes a vitellogenin-2-like protein.

### Differential expression of piRNA clusters in eggs and adults

Replicates separated by clutch in Eggs and the F_0_ maternal generation (PCA plots, Figure 1), with principal component 1 accounting for 60% of variation in eggs versus 29% in F_0_ adults. This was lost in F_1_ adults and F_3_ generations which were inter-mixed by clutch (Figure S1). There were 578 differentially expressed clusters with expression greater in 1^st^ clutch eggs and 686 such clusters in 5^th^ clutch eggs (Table 2 and File S2, DESeq2 output). Although a large number of differentially expressed loci in each clutch, the magnitude of difference was modest for the majority of clusters, with an average log_2_-fold change of 0.56 for significant egg clusters (File S2, log_2_-fold change column). Bias is increased towards 5^th^ clutch eggs when considering clusters of longer than 10 kb, at 86 with expression greater in 1^st^ clutch eggs and 213 in 5^th^ clutch eggs. Only 130 of all 1264 differentially expressed loci exhibited a doubling in expression in one clutch versus the other (log_2_-fold change greater than one or less than minus one), and the majority of these had low absolute expression levels. When restricting differentially expressed loci to those with a count of greater than 1000 transcripts per million (TPM) 41 and eight 1^st^ and 5^th^ clutch clusters were significant respectively. Of the eight 5^th^ clutch loci (Table S4), seven overlap a gene-body in the genome annotation, four of which are lncRNA loci. EggNOG annotated one cluster (NC_046177.1:4348409-4351253) as containing a Tudor domain (TDRD6) which has roles in germ cell development and the piRNA ping-pong cycle [36,37]. Related to the ping-pong cycle, three further clusters overlap ATP-binding helicase genes. However, none of these were differentially expressed in eggs or adults nor were they of high TPM expression. For the 41 clusters more highly expressed in the 1^st^ clutch, 22 overlapped predicted protein coding genes, 10 lncRNAs and two pseudogenes. Annotated protein-coding genes included Spermatogenesis-associated protein 20 (NC_046183.1:4720450-4720481), but no genes directly implicated in piRNA production or regulation. Additionally, 33 of the 41 1^st^ clutch clusters overlap a known transposable element versus zero for the 5^th^ clutch clusters, which was also reflected in 12 eggNOG annotations to repetitive element domains for 1^st^ clutch clusters.

**Table 2.** Differentially expressed piRNA clusters upregulated in each clutch relative to the other in each generation. TPM > 1000 = an equal to or greater than 1000 average transcripts per million threshold in the upregulated condition for egg comparisons.

| Comparison | Upregulated in 1<sup>st</sup> clutch | Upregulated in 5<sup>th</sup> clutch |
| --- | --- | --- |
| Eggs | 578 | 686 |
| Eggs, TPM > 1000 | 41 | 8 |
| F<sub>0</sub> | 179 | 170 |
| F<sub>1</sub> | 31 | 22 |
| F<sub>3</sub> | 0 | 0 |

**Figure 1.**
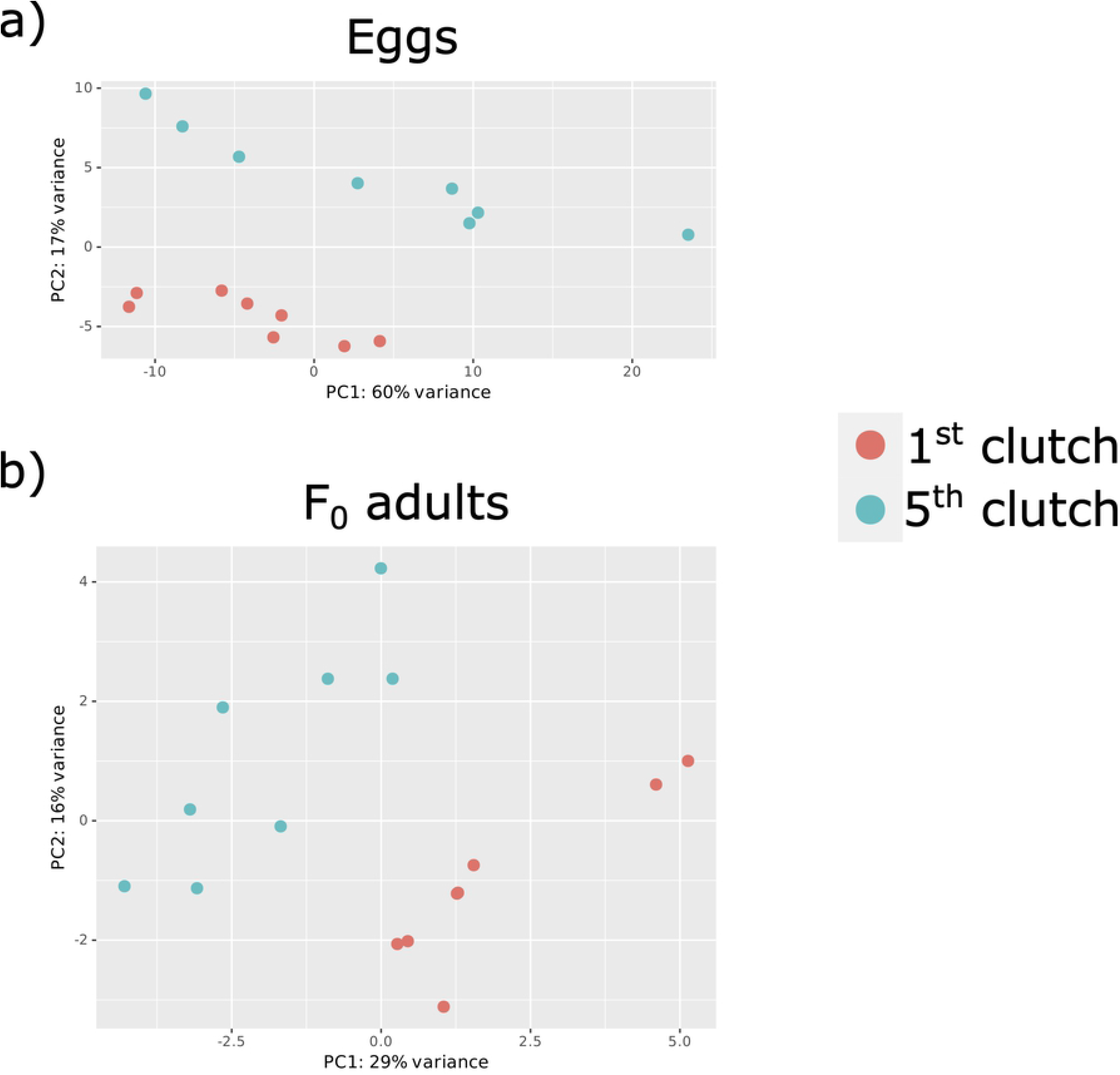
PCA plots of 1^st^ and 5^th^ clutch replicates for piRNA cluster expression of the 500 most variable clusters for (a) egg and (b) F_0_ generations.

In the maternal (F_0_) generation, 179 clusters were up-regulated in mothers on their 1^st^ clutch versus 170 in those on their 5^th^ clutches (Table 1). This dropped to 31 and 22 clusters in the F_1_ generation adults (whose mothers produced them when on either their 1st or 5^th^ clutch) and zero differences between clutches in the F_3_ generation (whose great grandmothers were on either their 1st or 5^th^ clutch). There was low concordance in cluster expression between adult and egg generations (Figure 2, Table S5), with less than 10% of egg clusters being shared with F_0_ adults and a smaller amount with F_1_ adults. This distinction between adult and egg libraries was supported by 14 of the highly-expressed proTRAC clusters being significantly differentially expressed in F_0_ adults. Whereas only one proTRAC cluster (NW_022654559.1:2-5240) was differentially expressed in eggs, in that case more highly in 1^st^ clutch eggs. Of the 14 significant F_0_ clusters, 13 were more highly expressed in mothers on their 5^th^ clutch than those on their 1^st^, however the single proTRAC cluster more highly expressed young mothers is the same as that for eggs and had the lowest adjusted p-value of the 14 (4.84 × 10^−13^). This cluster likely encodes a transposase according to its EggNOG annotation (NW_022654559.1:2-5240, Table S6).

**Figure 2.**
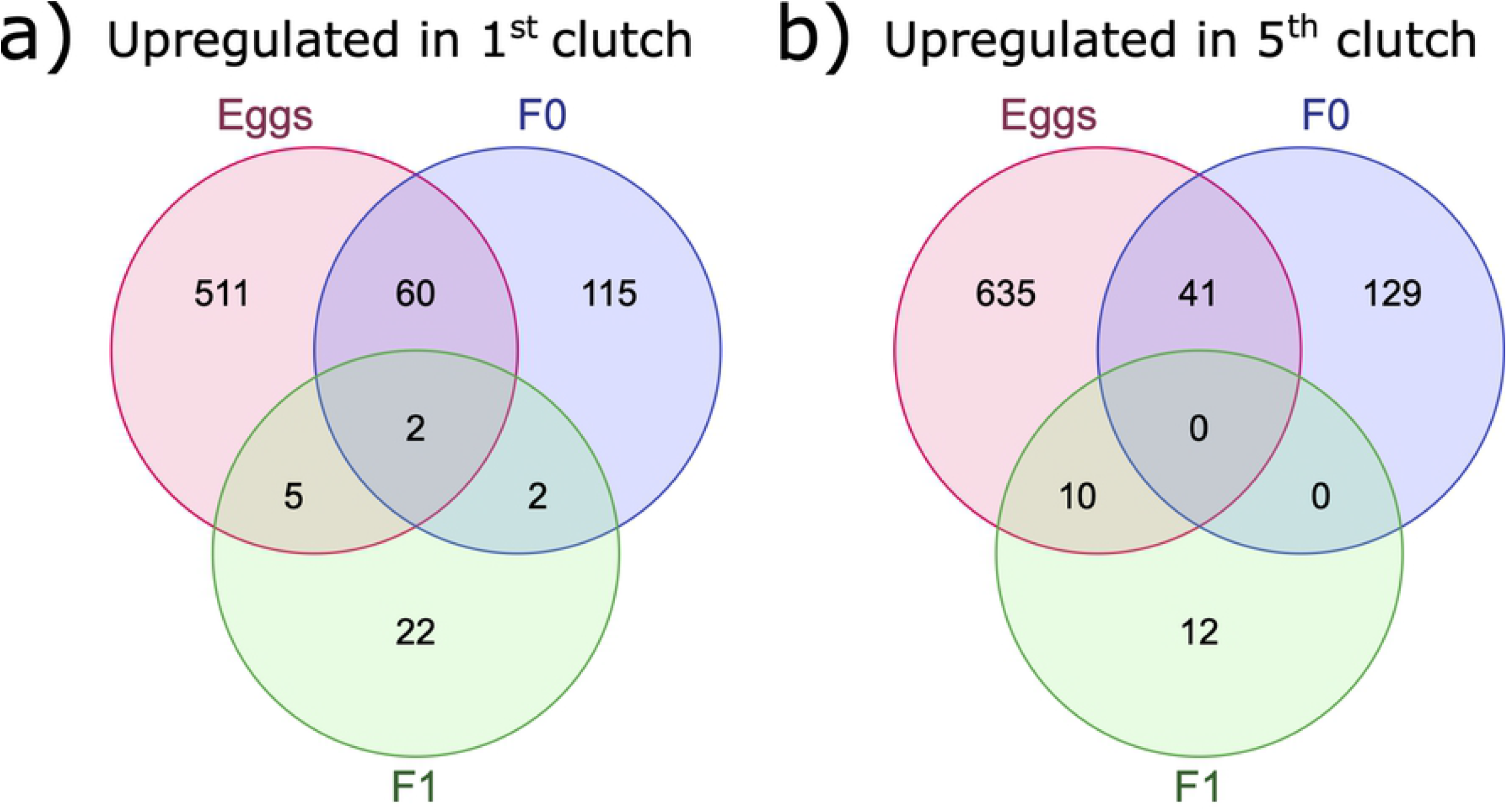
Venn diagrams of overlaps between eggs and F_0_ and F_1_ adult differentially expressed clusters. Part ‘a)’ shows clusters upregulated in the 1^st^ clutch versus 5^th^ clutch and ‘b)’ clusters upregulated in the 5^th^ clutch versus 1^st^ clutch.

## Discussion

By a series of filtering steps, a high-confidence and well-replicated piRNA dataset was created for 1^st^ and 5^th^ clutch eggs, their parents (the F_0_), and descendent generations of parthenogenetically reproducing *D. magna* adults (the F_1_ and F_3_ generations). By integrating two methods of predicting small RNA loci, piRNA clusters were then quantified and contrasted between clutches representing different ages of *D. magna*. A large number of clusters were differentially regulated in eggs, and much less so in whole adult tissues of each generation tested. Eggs and their parental F_0_s showed good separation of expression patterns (Figure 1), a result in line with miRNAs derived from the same dataset [32]. Hence, the piRNA profile of eggs is likely dictated by maternal provisioning of the eggs as in *Drosophila* [1,38,39]. This commonality was lost by adulthood, as in the F_1_ generation, adults were inter-mixed in expression profiles and differential expression was much reduced between clutches compared to F_0_s and eggs.

### *D. magna* piRNA cluster profiles

Despite covering only 2.1% of the piRNA producing loci, proTRAC-predicted clusters were responsible for approximately ∼50% of piRNAs. In this, *D. magna* is similar to *Drosophila* where such long clusters of fragmented transposons, which proTRAC was designed to detect, are also responsible for producing the majority of piRNAs [8,40]. Only mono-directionally expressed piRNA clusters were predicted by proTRAC in *D. magna* egg and adult tissues. Bi-directionally expressed piRNA clusters may represent a fly-specific adaptation [9,41], however a minority of piRNA clusters (15%) in the mud crab crustacean, *Scylla paramamosain*, were also expressed in this manner [42].

### *D. magna* parthenogenetically-reproducing piRNA clusters

All egg data surveyed was from embryos destined to become asexual females. The advantage of which is the genetic identity of replicates, which was expected to reduce variance in response to age. Conversely, this means males were not sampled in this study and testis are known to be a location of piRNA expression in arthropods [42–44]. Hence, this study is not an exhaustive survey of piRNAs in *D. magna*, as distinct piRNA clusters may well be expressed in male germlines and/or somatic tissues. In *S. paramamosain*, piRNA expression and cluster activity was much greater in ovaries than testes [42], and predicted piRNA clusters sizes were longer and more numerous in ovaries than testis. Indeed, although not an exact comparison, the number of expressed ovary piRNA clusters in *S. paramamosain* was much greater than that for *D. magna* eggs (19). Sensitivity to proTRAC sliding window parameters may contribute to these different estimates [45]. In [42], 300 bp windows with increments of 100 bp were applied versus 5000bp windows across 1000bp increments here. The phylogenetic divergence between *S. paramamosain* and *D. magna* may also play a role, as they belong to the Malacostraca and Branchipoda, respectively.

### Tudor domain genes regulated by piRNAs

Tudor domain genes are thought to form a molecular scaffold that connects elements of the ping-pong cycle [46]. A piRNA cluster containing such a domain (TDRD6) was upregulated in 5^th^ clutch eggs which suggests a self-regulating feedback loop may occur in the ping-pong cycle. If piRNAs are down-regulating a component of their own pathway in one treatment we would expect to see lower general expression of clusters relative to comparisons. This is indeed what we observed here with more clusters with a TPM > 1000 over-expressed in 1^st^ clutch versus TDRD6-upregulated 5^th^ clutch eggs at 41 and eight clusters respectively. This hypothesis is speculative and would require implication of the identified TDRD6 domain encoding gene in piRNA production in *Daphnia* and supporting gene expression data for 1^st^ and 5^th^ clutch eggs. Furthermore, 33 of 41 clusters with TPM > 1000 upregulated in 1^st^ clutches were annotated with a TE element or domain versus zero in 5^th^ clutches, suggestive of more efficient TE control in 1^st^ clutch parthenogenetically-reproducing *D. magna*. This is in line with *Drosophila* somatic tissues [19–21], but not with the idea of continued efficient policing of reproductive tissues with age [17,18,23]. No other differentially expressed piRNA clusters overlapped piRNA pathway genes. Three piRNA clusters of low TPM expression in eggs and adults did overlap ATP-binding helicase genes which act during oogenesis alongside piRNAs to methylate and repress transposable elements [47].

### PiRNA cluster versus piRNA targeting

We restricted this analysis to regions of the genome from which piRNAs originate. This is because most small RNA prediction algorithms have been designed to identify miRNA targets according to the well understood seed matching rules of miRNA-target interactions. Such methods are prone to false positives [48,49]. Because of this, Fridrich et al [48] recommend biological interpretation of miRNA interactions in combination with experimental result. Currently, it is hard to justify using such imprecise algorithms for piRNA interactions as they were not designed for this (but see [50–55] among others). Despite this, recent advances in understanding the seed-matching rules of piRNAs indicate longer and therefore more specific piRNA-target interactions [13,56,57], rules which appear to be shared with *Aedes* mosquitoes [58]. If a common property across animals, more limited potential targets per piRNA than miRNA will perhaps make computational inference of piRNA targets a more fruitful exercise than it has been for miRNAs.

## Conclusions

By strict and multi-step filtering it was possible to enrich a small RNA dataset for piRNAs. Clusters of transposable element fragments covered a small fraction of the total predicted piRNA-producing loci but were responsible for most piRNA expression. These clusters were all transcribed in a single direction. Differential expression results were interpreted by piRNA clusters versus targets due to false-positive issues with miRNA-based targeting approaches. Recent insights into piRNA seed-matching to targets, however, may make piRNAs more amenable than miRNAs to computational inference of targets in future. The dynamics of piRNA cluster expression in *Daphnia magna* changes with age in adults and their eggs, but not in the resulting adults or subsequent generations, as was observed for miRNAs in this *D. magna*. Our results suggest more efficient control of TEs in younger 1^st^ clutch *Daphnia* than in older *Daphnia* on their 5^th^ clutch, perhaps through a piRNA-mediated negative feedback on a Tudor domain containing components of the piRNA pathway, a hypothesis requiring further investigation.

## Methods

### Maternal ageing experiment and sequencing

This experiment was first described in Hearn et al 2018 [32] and is summarized here. A clone of *Daphnia magna* (C32) originating in Kaimes pond near Leitholm in the Scottish Borders, United Kingdom [59] shows a maternal effect pattern where large offspring are produced when mothers are older [60]. To create experimental lines, 24 groups of five female *Daphnia* were placed into jars of 200 ml of artificial culture medium and fed 2.5 × 10^6^ cells of the single-celled green algae *Chlorella vulgaris* daily for three generations. After three generations, five second clutch new-born females were designated F_0_ of the experiment and fed *ad libitum* (5 × 10^6^ cells *C. vulgaris* daily). Eggs of the 1^st^ and 5^th^ clutch of these acclimatized F_0_ mothers were collected (Figure 3a). Eggs were collected by flushing brood chambers with medium using a hypodermic syringe, pipetted onto tissue paper to dry, and then ground by motorized pestle in 350 µl Qiazol. Eggs from six jars each containing five *Daphnia* were combined to form a biological replicate from 30 *Daphnia* mothers in total. Eight biological replicates from each 1^st^ and 5^th^ clutch mother were created resulting in 16 egg libraries in total.

**Figure 3.**
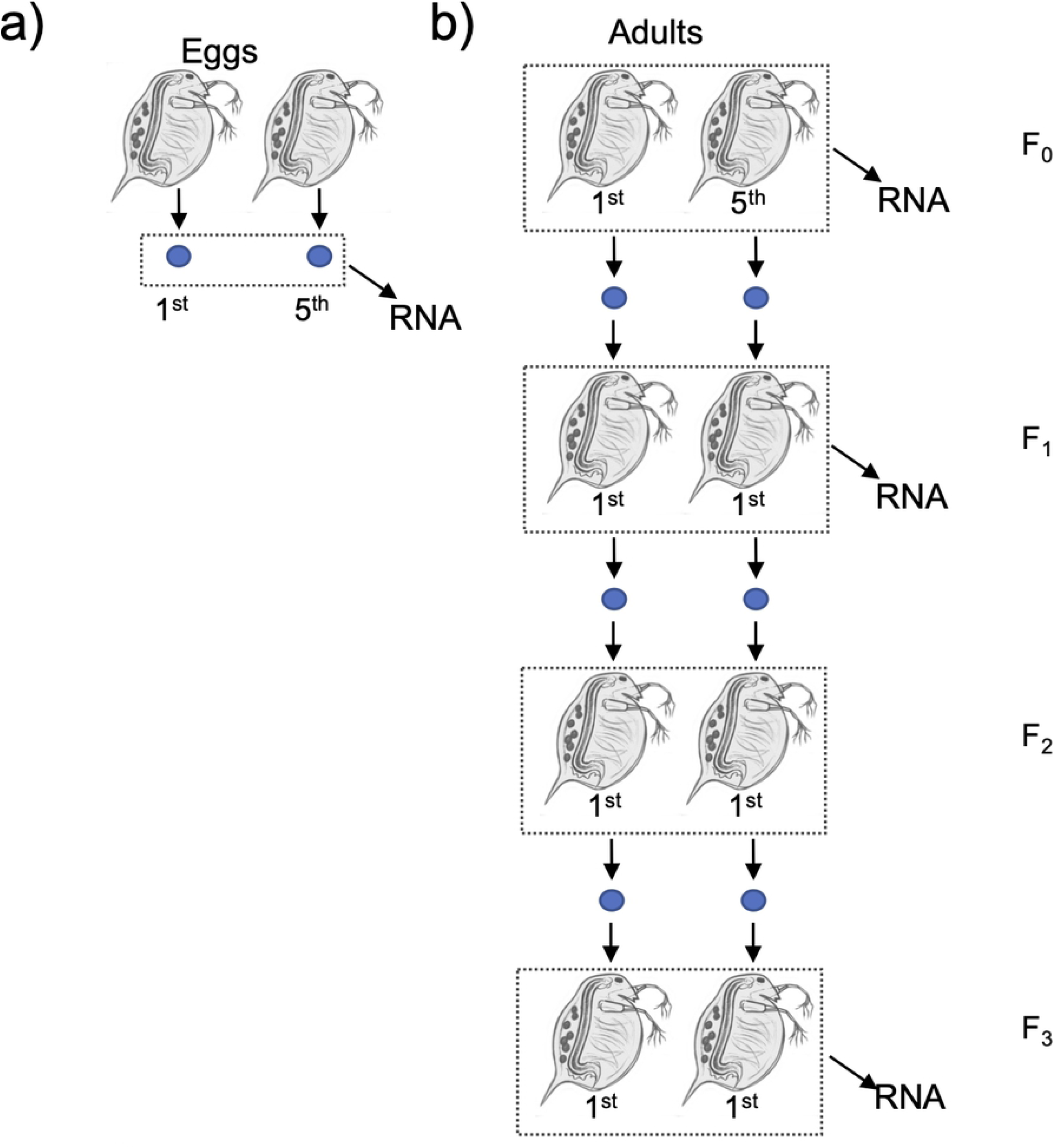
Schematic showing design for experiments to generate sRNA pools. **a) Eggs and b) adults of different ages (F_0_) or whose parents or great grandparents were of different ages (F_1_ and F_3_)**. Numbers below Daphnia mothers/eggs indicate the clutch sampled from for that generation. Adapted from Hearn et al 2018, Figure 2.

Adult reference libraries were created separately from eggs (Figure 3b). First, replicate young (i.e., on their first clutch) and replicate old mothers (on their fifth clutch) were harvested to produce the F_0_ set of RNA samples. Prior to RNA harvesting, new-born from these young and old F_0_ mothers were isolated and grown to adulthood. This next generation of adults constituted the F_1_, and RNA from F_1_ adults was harvested just after they had their first clutch, the idea being to test for a maternal effect of having been born to a young or old (F_0_) mother [32]. The first clutch offspring of F_1_ adults were used to seed an F_2_ generation, from which RNA was not harvested. The first clutch offspring of F_2_ adults were used to seed an F_3_ generation, from which we harvested RNA when they had their first clutch, in order to test for great-grandmaternal effects, where great grandmothers (i.e., the F_0_) were either young or old when they produced eggs. Each replicate consisted of five adults per jar and there were eight replicates per treatment per generation. All generations were fed *ad libitum* throughout the experiment.

In total, 64 single-end small RNA libraries of 50 bp length were prepared with the CleanTag Small RNA Library kit (16 each F_0_ adults, F_0_ eggs, F_1_ adults, and F_3_ adults) and sequenced at Edinburgh Genomics (Edinburgh, United Kingdom) to 50 bp length [32]. Raw sequence data were deposited under bioproject PRJEB22591 in the European Nucleotide Archive.

### Identification of *D. magna* piRNAs and differential expression analysis

Raw reads were trimmed with fastp (v0.20.1) under defaults and reads 21 bp longer and 35 bp or under retained and converted to fasta using seqret (EMBOSS v6.5.7.0). Reads were aligned to all *Daphnia* sequences in the RFAM database (release 14.5) and to pre-miRNA sequences identified in [32,61] using SortMeRNA (v4.3.3). The filtered reads were classified as piRNA or not in piRNN using the *Drosophila melanogaster* trained model [62]. This classifier is based on a convolutional neural network framework and was recommended for use with none model organisms after comparison with other classifiers [45].

Two approaches were combined to identify piRNA producing loci across the *Daphnia magna* chromosomal genome assembly [25] using reads classified as piRNAs (Figure 4, piRNA filtering flow diagram). Firstly, piRNA producing clusters were identified for adult (F_0_ + F_1_ + F_3_) and egg (F_0_ egg) libraries separately in proTRAC (v2.4.2) [63]. The proTRAC pipeline removes collapses redundant reads (“TBr2_collapse.pl”), removes low-complexity sequences (“TBr2_duster.pl”), aligns reads to the genome (using “sRNAmapper.pl”) and reallocates multi-mapping reads according to local transcription levels (“reallocate.pl”) before the proTRAC algorithm itself is run. Each step was run with the proTRAC v2.4.2 documentation example settings. The second approach was to align the piRNA-classified reads to the genome using the small RNA aligner ShortStack (v3.8.5) [64] allowing the maximum two mismatches per mapping to account for differences between the genome assembly and strain C32. Each ShortStack-defined cluster was checked for the characteristic 10 bp 1U and 10A overlap (the ping-pong signature) in read-alignment stacks with in PingPongPro (v1.0), and any such sequences combined with others if within a range of 1000 bp of one another (option “T 1000”). ShortStack clusters with significant ping-pong signals (adjusted p-value <-0.05) were combined with the Adult and Egg proTRAC predictions using bedtools merge (v2.23.0). Reads were then re-aligned to the genome in ShortStack using the combined piRNA cluster co-ordinates to quantify each cluster per library for differential expression analysis. Finally, clusters were removed if they overlapped another species of RNA in the *Daphnia magna* genome annotation (defined as “guide_RNA”, “rRNA”, “snoRNA”, and “snRNA” in the assembly annotation file “GCF_003990815.1_ASM399081v1_genomic.gff”) and, to avoid undetected miRNAs, if the average read length aligned to a cluster was less than 24 bp. Differential gene expression between 1^st^ and 5^th^ clutches was performed on counts per cluster per library separately for eggs and adult generations in the R Bioconductor package DESeq2 (v1.32.0) [65]. P-values were adjusted using Independent Hypothesis Weighting [66] in the R Bioconductor package IHW (1.20.0) as part of the DESeq2 workflow, and a significance threshold for adjusted p-values of < 0.05 applied. For comparison of expression between piRNA clusters, counts were converted into transcripts per million (TPM) values using “counts_to_TPM.R” (https://gist.github.com/slowkow/c6ab0348747f86e2748b). Mean fragment length per cluster used to calculate TPM was the average read length of reads aligned to that cluster. Unlike most single-end sequencing experiments, the length of the molecule of cDNA sequenced (fragment length) corresponds to the RNA sequenced and was shorter than the read-length sequenced (50 bp), in this case putative piRNAs of approximately 26 bp. Differences in TPM values averaged across replicates were used to compare egg and combined (F0, F1 and F3) adult differences to identify tissue-biased piRNA clusters. It was not appropriate to perform a DESeq2 analysis due to the distinct and time-separated library preparation between eggs and adults undertaken in [32]. A flow chart of bioinformatic steps from raw-reads to DESeq2 differential expression testing was created using https://app.diagrams.net/. Venn diagrams intersecting genes upregulated in 1^st^ and 5^th^ clutches between eggs, F_0_ and F_1_ adults were created using https://www.molbiotools.com/listcompare.php. R code for differential expression analyses and TPM count generation is given in File S3.

**Figure 4.**
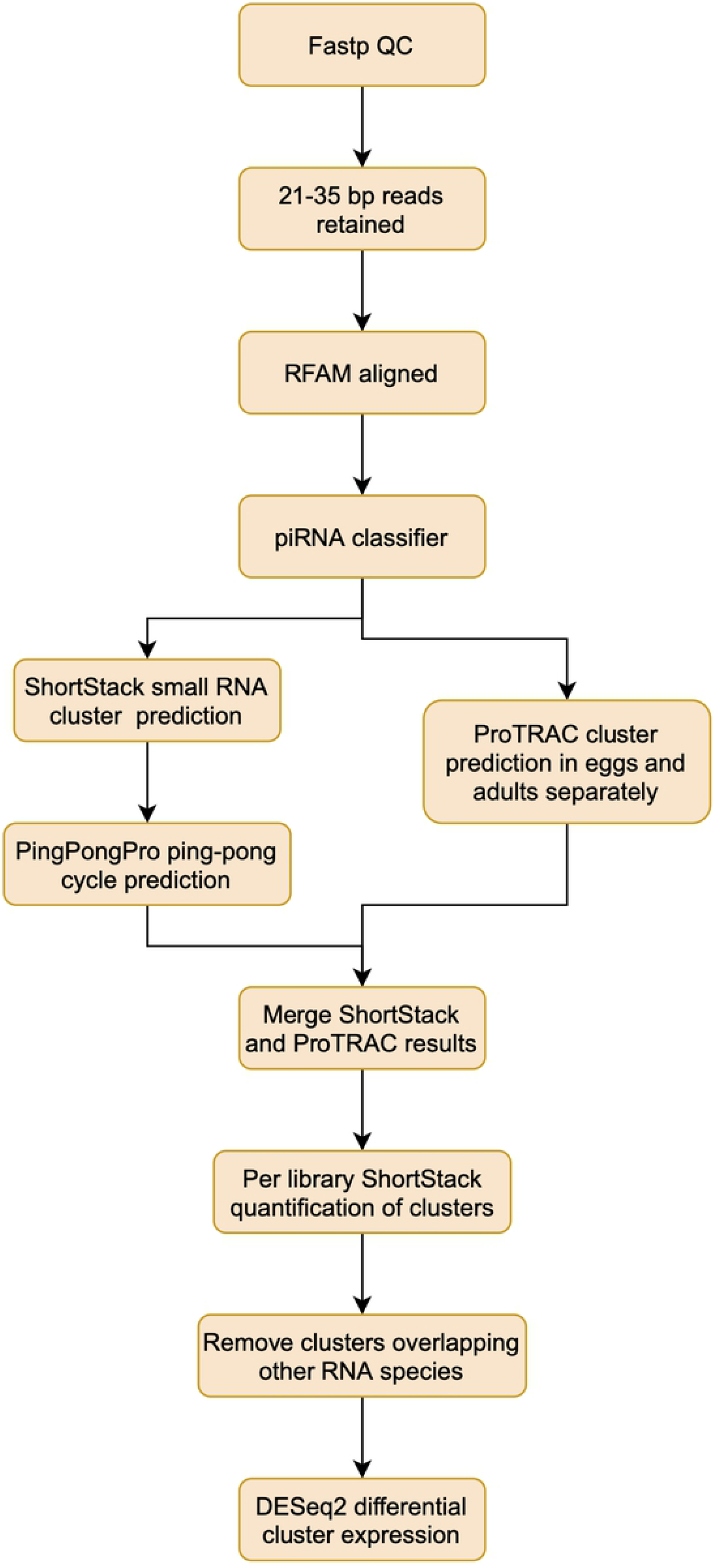
Flow diagram of piRNA filtering, cluster prediction and quantification steps.

### Annotating piRNA clusters

In order to identify piRNA regulated transposable elements, the *D. magna* genome was annotated with RepeatModeler (v2.0.2a) and RepeatMasker (4.1.2) [67,68] using predicted repeats and Dfam database transposable elements (version 02/09/2020). The Repeatmasker gff file was then filtered to remove simple and low-complexity repeats prior to intersection with piRNA cluster locations. PiRNA clusters were also annotated independently using eggNOG-mapper.

## Acknowledgements

This research was funded by Leverhulme Trust grant RPG-2015-406 and Institutional Strategic Support Funds (204804/Z/16/Z) awarded to School of Biological Sciences, University of Edinburgh by The Wellcome Trust.

## Supplementary Tables

**Table S1. Raw and filtered read counts per library**. For raw, fastp, Rfam and piRNN classifier steps with percentages remaining after filtering step and of original raw reads for the final piRNN step.

**Table S2. Total and proTRAC-predicted piRNA cluster normalised read counts per library**. For all replicates included in experiment, percentage of proTRAC cluster reads are given in the “percent proTRAC” column.

**Table S3. PiRNA clusters with transcript per million (TPM) differences greater than 1000 between egg and adult libraries**. Egg TPMs were the average of 1^st^ and 5^th^ clutch replicates combined and adult TPMs were the average of all adult F_0_, F_1_ and F_3_, 1^st^ and 5^th^ clutches combined.

**Table S4. PiRNA clusters differentially expressed in eggs with average transcript per million (TPM) counts greater than 1000 in the upregulated replicates**.

**Table S5. Overlap between clusters by F**_**0**_, **egg and F**_**1**_ **generations in each direction**. Overlap is the number of shared differentially expressed clusters in a concordant direction (for example, both upregulated in 1^st^ clutch) between all eggs, F_0_ and F_1_ generations.

**Table S6. EggNOG annotations for clusters included in the experiment**. Clusters not included in this file were not annotatable with EggNOG.

## Supplementary Figures

**Figure S1. PCA plots of 1**^**st**^ **and 5**^**th**^ **clutch replicates for piRNA cluster expression of the 500 most variable clusters for (a) F**_**1**_ **and (b) F**_**3**_ **generations**.

## Supplementary Files

**File S1. ProTRAC piRNA cluster predictions for egg and adult libraries**.

**File S2. Significantly differentially expressed piRNA results generated in DESeq2 for each generation 1**^**st**^ **versus 5**^**th**^ **clutch comparison**. Four intra-generation comparisons were made: egg, F_0_, F_1_ and F_3_. There were zero differentially expressed F3 clusters hence only three sets (egg, F_0_ and F_1_) of DESeq2 results are listed.

**File S3. DESeq2 R code for calculating differential expression between 1**^**st**^ **(labelled “Young”) and 5**^**th**^ **(“Old”) for each generation (eggs, F**_**0**_, **F**_**1**_ **and F**_**3**_**) given consecutively, TPM values and PCA figures**.

